# Novel Data Transformations for RNA-seq Data Analysis

**DOI:** 10.1101/432690

**Authors:** Zeyu Zhang, Danyang Yu, Minseok Seo, Craig P. Hersh, Scott T. Weiss, Weiliang Qiu

## Abstract

We propose eight data transformations for RNA-seq data analysis aiming to make the transformed sample mean to be representative of the distribution center since it is not always possible to transform count data to satisfy the normality assumption. Simulation studies showed that limma based on transformed data by using the *rv* transformation (denoted as limma+*rv*) performed best compared with limma based on transformed data by using other transformation methods in term of high accuracy and low FNR, while keeping FDR at the nominal level. For large sample size, limma based on transformed data by using the 8 proposed transformation methods had similar performance to limma based on transformed data by using existing transformation methods for equal library size scenarios. Otherwise, limma based on transformed data by using the *rv*, *lv*, r*v2*, or *lv2* transformation, or by using the existing *voom* transformation performed better than limma based on data from other transformation methods. Real data analysis results showed that limma+ *l2* performed best, while limma+ *rv* also had good performance.

## Background

With the rapid development of next-generation high throughput RNA sequencing technologies in recent years, genomics studies have seen tremendous advancement. RNA-seq technology is a type of next generation sequencing technology to estimate the expression levels of genes in whole-genome scale studies and has become the standard technology for the study of genomics [1, 2]. RNA-seq technology can help identify new genes, with high-sensitivity, high signal-to-noise ratio and small sample requirements. Also, RNA-seq technology can measure read counts at exons, genes, or gene units. Therefore, RNA-seq sequencing technology has been widely used in many different research fields[3, 4].

RNA-seq data are usually represented by a matrix of counts showing the expression levels of mRNAs (rows) for a set of samples (columns) after processes such as adapter remove step, alignment step, and quantification step. For each sample, millions of reads can be measured by the RNA-seq technique[5]. According to the gene annotation and genome build, numbers of features might be different. Different pipelines, such as Cufflink pipeline, Hisat2-StringTie pipeline, and Star-FeatureCount pipeline could result in different properties of the count matrix. Two common properties are sparsity and skewness. Sparsity means that many counts in the count matrix are zero. Skewness means that the histogram of all counts in the count matrix is usually skewed. Skewness indicates that data transformation is required to apply linear regression analysis, which assumes data from normal distributions. Sparsity indicates that the log2 transformation, which is commonly used in gene microarray data, could not be directly applied to RNA-seq data analysis since log2(0) does not exist. It is still expensive to collect RNA-seq data for large sample size. Hence, existing RNA-seq datasets usually have small sample size. To address these two common properties, count distributions, such as Poisson, negative binomial, and inflated Poisson distributions, have been proposed to fit RNA-seq data[6, 7]. Commonly used R Bioconductor packages that fit RNA-seq data using count distributions include edgeR[8, 9], DESeq[10], and DESeq2[11]. These methods could borrow information across genes to increase the power of the tests for detecting genes differentially expressed between two conditions (e.g., cases versus controls).

The distributions of counts are not as statistical tractable as normal distributions[12]. Moreover, there are much fewer analytic tools for count distributions than there are for normal distributions in statistical analysis. Law et al. (2014)[12] proposed the *voom* transformation to transform the count distribution to a distribution close to the normal distribution in RNA-seq data analysis and demonstrated that using *limma*[13] with *voom*-transformed count data performed comparable to count-based RNA-seq analysis methods, such as edgeR[8, 9], DESeq[10], baySeq[14] and DSS[15].

The *voom* transformation is a sample-specific transformation, defined as log-counts per million (log-cpm):

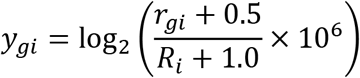

where r_gi_ is the count of the g-th mRNA gene for the i-th sample, R_i_ is the total counts (R_i_=r_1i_+r_2i_+…+r_Gi_) for the i-th sample, g=1, …, G, i=1, …, n, G is the number of mRNA genes, and n is the number of samples.

The goal of the *voom* transformation is to make the empirical distribution of transformed RNA-seq data closer to a normal distribution so that the moderate t tests (*limma*) could be used. However, it is not always possible to transform count data to have a distribution closer to a normal distribution in real data analysis[16]. For example, for the SEQC data, that was analyzed in real data analyses part of [12], the empirical distribution (i.e., histogram) of the *voom* transformed data is still far from a normal distribution (Figure 1). The histogram is based on the pooled data Ygi, g=1, …, 92, i=1, …, 8.

**Figure 1.**
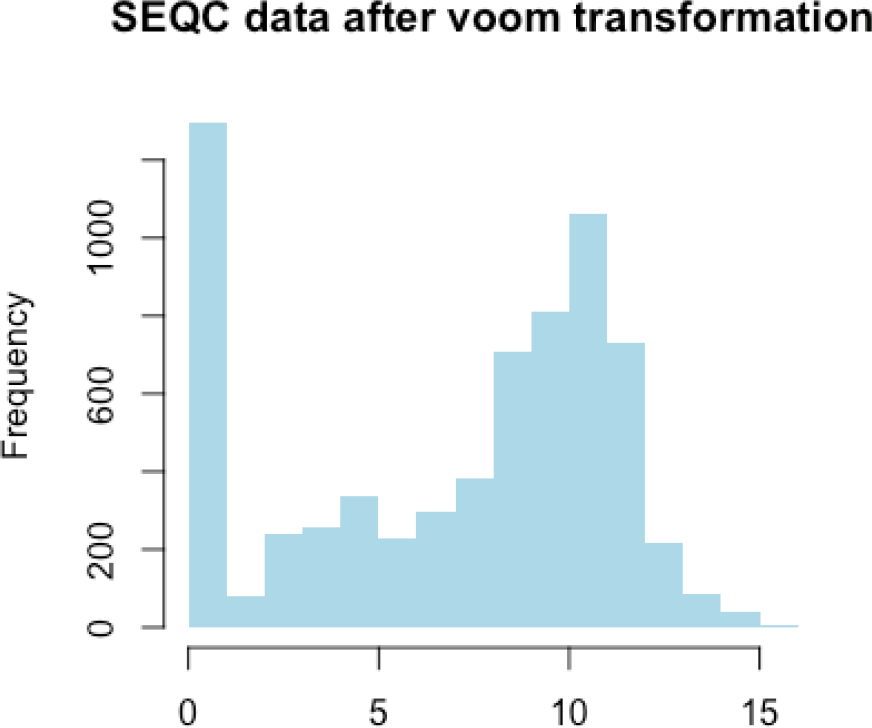
Histogram of the pooled SEQC RNA-seq data after the *voom* transformation. The histogram showed that the empirical distribution after the TMM scale normalization, quantile normalization and *voom*-transformation could still be far from a normal distribution.

In this article, we proposed to relax the normality requirement for a data transformation. Most statistical models, such as two-sample t-test, focus on the centers of the two distributions to check if two distributions are same or not. Sample means are usually used to represent distribution centers. However, for skewed distributions, sample means are not good to characterize the distribution centers. Instead, sample medians are usually used to characterize the centers of skewed distributions. However, sample medians do not have as tractable properties as sample means. For instance, it is hard to derive the distribution of sample median. In this article, we aim to transform the RNA-seq count data by minimizing the difference between sample mean and sample median so that the sample mean would be a good representative to the center of the transformed distribution. Hence, most existing statistical models based on sample means, e.g., *limma*, can be directly applied to analyze transformed RNA-seq data.

## Results

### Results for simulation studies

In this article, we proposed 8 data transformation methods to improve the *voom* transformation. Four proposed transformations (*r*, *rv*, *r2*, and *rv2*) are based on root transformations. The other 4 proposed transformations (*l*, *lv*, *l2*, and *lv2*) are based on log transformations. To evaluate the effects of sample size and sample library on the performances of limma with data transformed by each of the 8 proposed data transformations and to compare them with the performance of limma with data transformed by the *voom* transformation, we performed eight simulation studies based on the simulation scheme in [12]. For each simulated RNA-seq dataset, we applied *limma* to the transformed data.

In addition, we would like to evaluate if using non-parametric approaches would have better performance than using parametric approaches in analyzing RNA-seq data, the distribution of which is non-normal. Specifically, we used the *Wilcoxon rank sum* test (denoted it as *Wilcoxon*) on each gene based on the *untransformed* counts. We then adjusted p-values to control false discovery rate < 0.05.

Figure 2 (a) shows that for scenarios with small sample size and equal library size (nCases=nControls= 3), limma with all the eight proposed transformations performed better than limma with the *voom* transformation in terms of *accuracy*. The accuracy is defined as the proportion of agreement between the true gene significance and the detected gene significance. Gene significance indicates if the gene is differentially expressed between cases and controls. The detected gene significance of a gene is 1 if the false discovery rate (FDR) adjusted p-value for this gene is < 0.05; the significance is 0 otherwise. Especially, limma with *l*, *rv*, *lv*, *l2*, *rv2* and *lv2* performed much better than *Wilcoxon* approach and limma with *voom*. Figure 2 (b) shows that for small sample size and equal library sizes scenarios, limma with all the eight proposed transformations had median FDR values ≤ 0.05, indicating that the proposed methods can control multiple testing well. Figure 2 (c) showed that limma with all the 8 proposed transformations had much lower median FNR than *Wilcoxon* approach and limma with *voom*, indicating that the proposed methods had much higher testing power than *Wilcoxon* approach and limma with *voom* when the sample size is small.

**Fig. 2.**
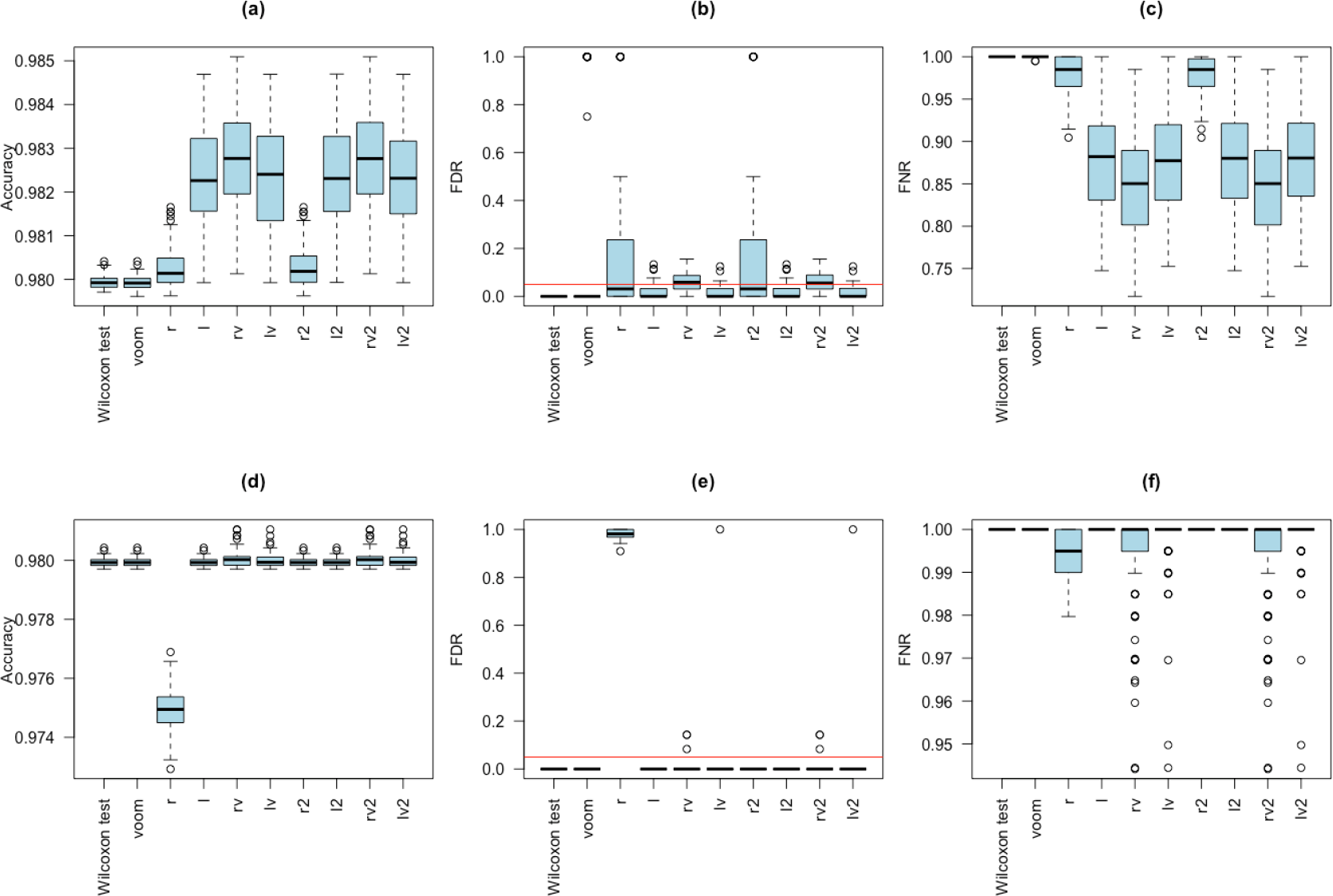
Results based on the 100 simulated datasets with small sample size (nCases=nControls=3). Upper pannel **(a,b,c)**: samples having equal library size; Lower panel **(d,e,f)**: samples having unequal library sizes.

Figure 2 (d) showed that when samples have different RNA-seq library sizes, limma with *rv* and limma with *rv2* had higher accuracy than limma with *voom*, while limma with the other 6 proposed transformations had similar performance to limma with *voom,* except for limma with the *r* transformation. Also, all the 10 methods we compared, except for limma with the *r* transformation, have very good performance in terms of FDR (Figure 2 (e)). All methods, except for limma with *r*, *rv*, and *rv2*, had median FNR equal to one (i.e., all true positives were identified as negatives), indicating all the methods had lower testing power when sample size is small and samples have different library sizes.

We also did simulations with large sample size (nCases=nControls=100) to compare the performance of the 10 different methods. Supplementary Figure 1(a) showed that when library sizes are equal, the methods (limma with *r*, *rv*, *r2*, and *rv2*), which are based on the root transformations, had slightly higher median accuracy than *Wilcoxon* and limma with *voom*. Supplementary Figure 1(b) showed that all 10 methods had mean FDR ≤ 0.05. Supplementary Figure 1(c) showed that all 10 methods had low median FNRs, which are lower than 0.015. Overall, the 10 methods had similar performance when sample size is large and sample library sizes are equal.

Surprisingly, the *Wilcoxon* test without data transformation performed best in terms of accuracy (Supplementary Figure 1 (d)) by using Law et al.’s (2014)[12] simulation studies when the sample size was large and samples had unequal library sizes. *Wilcoxon* also had the lowest FDR (Supplementary Figure 1 (e)) and low FNR (Supplementary Figure 1 (d)). However, *Wilcoxon* had significant higher FNR than limma with *voom*, *rv*, *lv*, *rv2*, and *lv2* (Supplementary Figure 1 (f)), indicating that *Wilcoxon* had lower power than limma with *voom*, *rv*, *lv*, *rv2*, and *lv2*.

Supplementary Figure 1 (d) showed that limma with *l*, *r2,* and *l2* had significantly higher accuracy than limma with *voom* and showed that limma with *r*, *rv*, *lv*, *rv2*, and *lv2* had slightly lower accuracy than limma with *voom*. Supplementary Figure 1(e) showed that all 10 methods had median FDR ≤0.05 and showed that *Wilcoxon* and limma with *r*, *l*, *r2*, and *l2* had much lower FDR than limma with *voom*. Supplementary Figure 1(f) showed that limma with *voom*, *rv,* and *rv2* had similar FNR and they had lower median FNR than other 7 methods.

Overall, the *Wilcoxon* test without data transformation can be used to detect differentially expressed genes for RNA-seq data when sample size is large and sample library sizes are unequal. If data transformation is preferred, then limma with *voom*, *rv*, or *rv2* can be used.

We also did simulation studies with total sample sizes 60 and 100 (i.e., nCases=nControls=30 and nCases=nControls=50). The results showed that when the total numbers of subjects is 60 or 100, *Wilcoxon* has the highest accuracy when samples had equal library size. Limma with the proposed transformations have similar accuracy to limma with *voom* when samples had equal library size (Supplementary Figure 2(a), Supplementary Figure 3(a)). When sample had unequal library sizes, limma with *voom*, *rv*, *lv*, *rv2* and *lv2* had similar accuracy to each other, and had a higher accuracy than *Wilcoxon* and limma with *r*, *l*, *r2*, *and l2* (Supplementary Figure 2(d), Supplementary Figure 3(d)). Supplementary Figure 2 (b, c, e, f) and Supplementary Figure 3 (b, c, e, f) showed that when equal library size, all 10 methods had median FDR ≤ 0.05 or close to 0.05, and that when unequal library size, limma with *voom*, *rv*, *lv*, *rv2* and *lv2 had* median FDR close to 0.05 and median FNR close to 0, indicating good error rates control.

Figure 3 and Supplementary Table 1 summarize the results of the eight simulation scenarios. For each simulation scenario, we used one-sample t-test to test the null hypothesis *H*_0_ that the mean of the 100 estimated FDR from the 100 simulated datasets is significantly ≤ 0.05 versus the alternative hypothesis *H*_a_ that the mean of the 100 estimated FDR from the 100 simulated datasets is significantly > 0.05. If the p-value of the one-sample t-test < 0.05, we reject *H*_0_. We counted the number of rejections for each method. For a given scenario, we also ranked median accuracy for methods that did not reject the null hypothesis *H*_0_. For ranks with ties, average ranks were used. For the methods that rejected the null hypothesis *H*_0_, we set their ranks as missing values.

**Figure 3.**
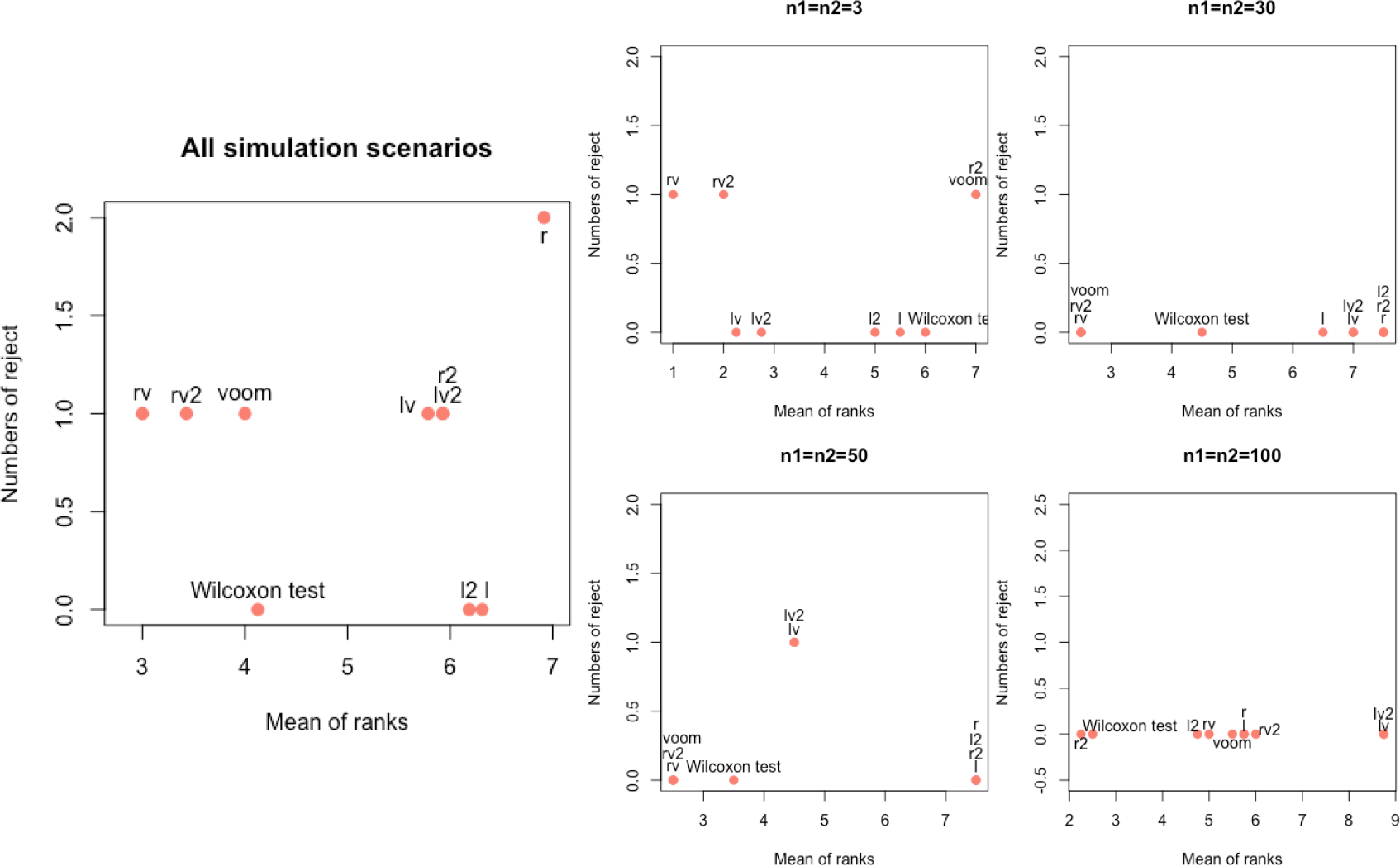
Summary of the performance of *Wilcoxon* and limma with all 9 transfromation methods in simulation studies.

The left panel of Figure 3 is the plot of the number of rejections versus mean rank considering all 8 scenarios and showed that limma with *rv* had the best performance with high accuracy, while keeping nominal FDR (0.05) in all simulation scenarios. We also can see that (1) limma with *rv2* is the second best; (2) *Wilcoxon* and limma with *voom* had the similar performance; (3) limma with *l* and *l2* kept nominal FDR (0.05) for all 8 scenarios, however their powers are low; and (4) limma with the *r* transformation had high false positive rate and low power. The right panel showed the plots for each sample size. We can see that limma with *voom* performed poorly for datasets with small sample size (nCases=nControls=3) or large sample size (nCases=nControls=100), while it is among the best methods for datasets with medium sample sizes (nCases=nControls=30 and nCases=nControls=50). Limma with *rv* and *rv2* performed best when sample size is relatively small (nCases=nControls < 100). For large sample size (nCases=nControls=100), limma with *r2* performed best and *Wilcoxon* test performed second, while all 10 methods kept a nominal FDR (0.05).

As we mentioned in the Background section, we aimed to transform the RNA-seq count data so that the sample mean would be representative of the center of the transformed empirical distribution. Hence, the difference between sample mean and sample median after transformation is an important judging criterion of RNA-seq transformation methods. We used the difference between sample mean and sample median based on the pooled expression levels of all genes and all samples to check if sample mean is close to the sample median. The smaller the difference, the closer the sample mean is to the distribution center. Figure 4 (a) and Figure 4 (c) showed that when sample library sizes are equal, all the 8 proposed transformations had the difference between sample mean and sample median very close to zero, while *voom* had relatively large mean-median differences. When sample library sizes are un-equal (Figure 4 (b) and Figure 4 (d)), *r*, *l*, *rv*, *lv*, *rv2*, and *lv2* still had the mean-median difference close to zero. However, *r2* and *l2* had much larger mean-median difference than zero. While *l2* had smaller difference than *voom*, the *r2* transformation had much larger mean-median difference than *voom*.

**Figure 4.**
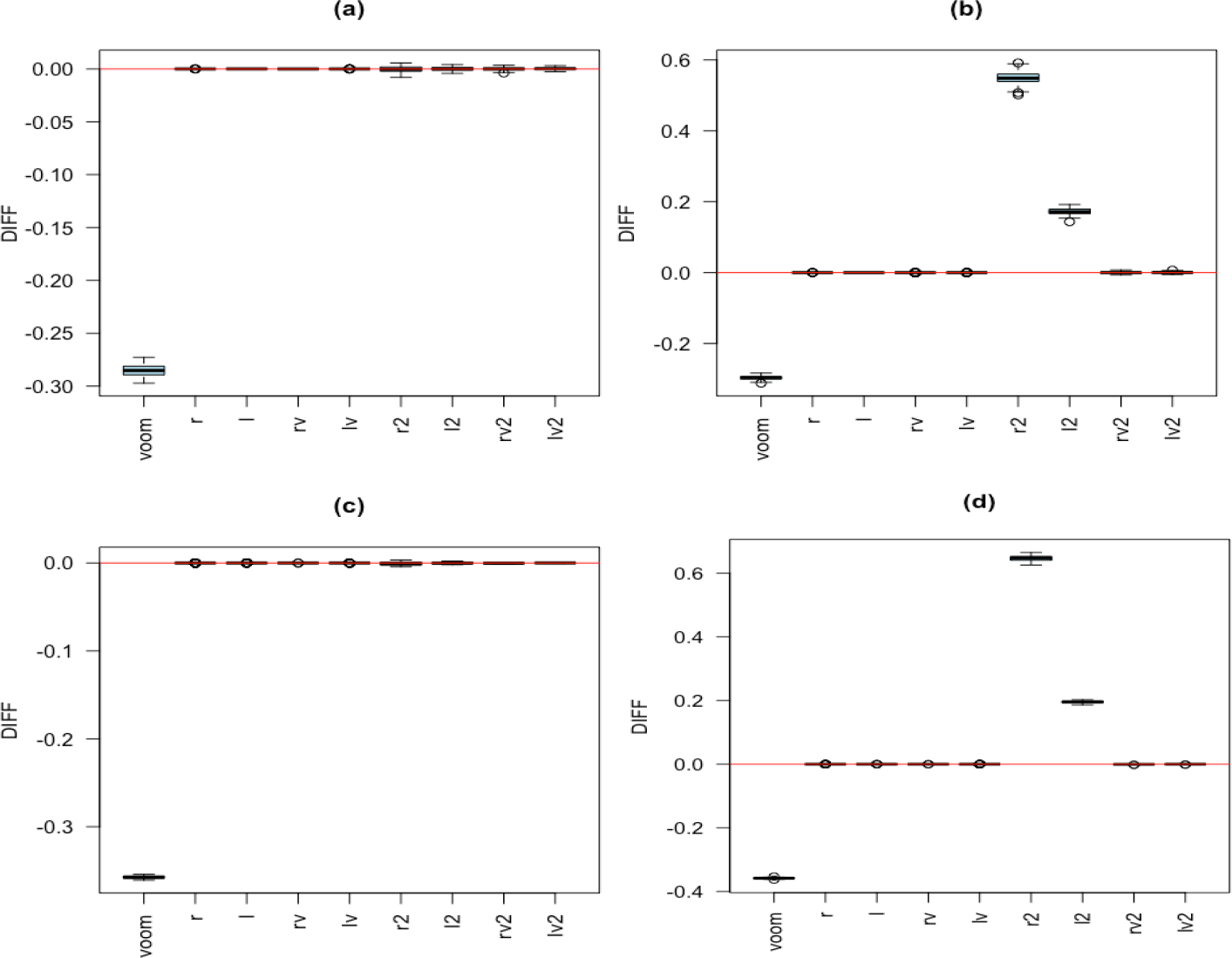
The difference (y-axis) between sample mean and sample median of the pooled expression levels of all genes and all samples after data transformation in four simulation studies. Lower panel: nCases=nControls=100. Left panel: samples with equal library size. Right panel: samples with un-equal library sizes.

### Real data analysis

The first two of our real data analysis datasets were based on the SEQC datasets[17]. We applied the *Wilcoxon* test, *limma* after the TMM scale normalization[18], quantile normalization, and the *voom* transformation (We still denoted the method as limma with *voom*), and *limma* after the 8 proposed transformations to the SEQC RNA-seq dataset.

The information about which genes are truly differentially expressed between two groups were determined based on qRT-PCR (Quantitative Real-Time PCR) experimental data. Figure 5 showed that limma with *r*, *l*, *rv, r2*, *l2, rv2* had higher accuracy than limma with *voom* and showed that limma with *lv* and *lv2* had equal accuracies to limma with *voom*. We noticed that *Wilcoxon*, a non-parametric method, failed to detect any true positives, although *Wilcoxon* had slighly higher accuracy to limma with *voom*.

**Figure 5.**
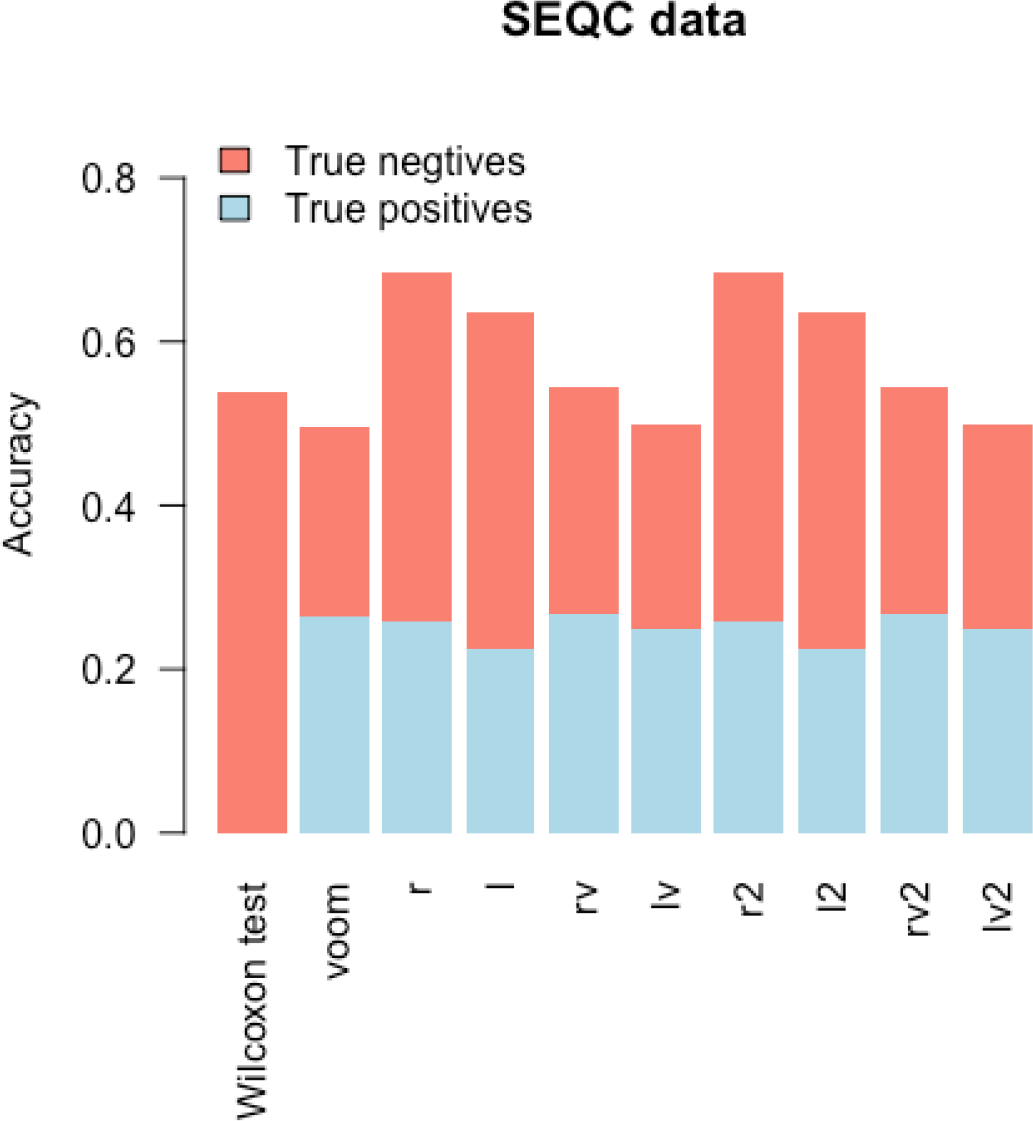
Accuracy for the SEQC dataset. The accuracy is plotted for each method and is split into true positive rate and true negative rate. Limma with the *r* and *r2* transformations had the highest accuracy (0.6836)

The analysis of the ERCC dataset is consistent with the SEQC analysis. The information about which genes are truly differentially expressed between two groups were determined based on concentrations of mixes (see Method Section). Figure 6 showed that limma with the *r*, *l*, *rv*, *r2*, *l2*, and *rv2* transformations had better accuracies than limma with *voom* and showed that limma with the *lv* and *lv2* transformations had equal accuracies to limma with *voom*. Specifically, limma with *r*, *l*, *rv*, *r2*, *l2*, and *rv2* had more true positives than limma with *voom*, indicating good testing powers. As in the analysis in the SEQC data, the *Wilcoxon* test failed to detect any true positives and had the lowest accuracy, which is consistent with the results of the simulation studies, indicating the *Wilcoxon* test has poor performance in datasets with small sample sizes.

**Figure 6.**
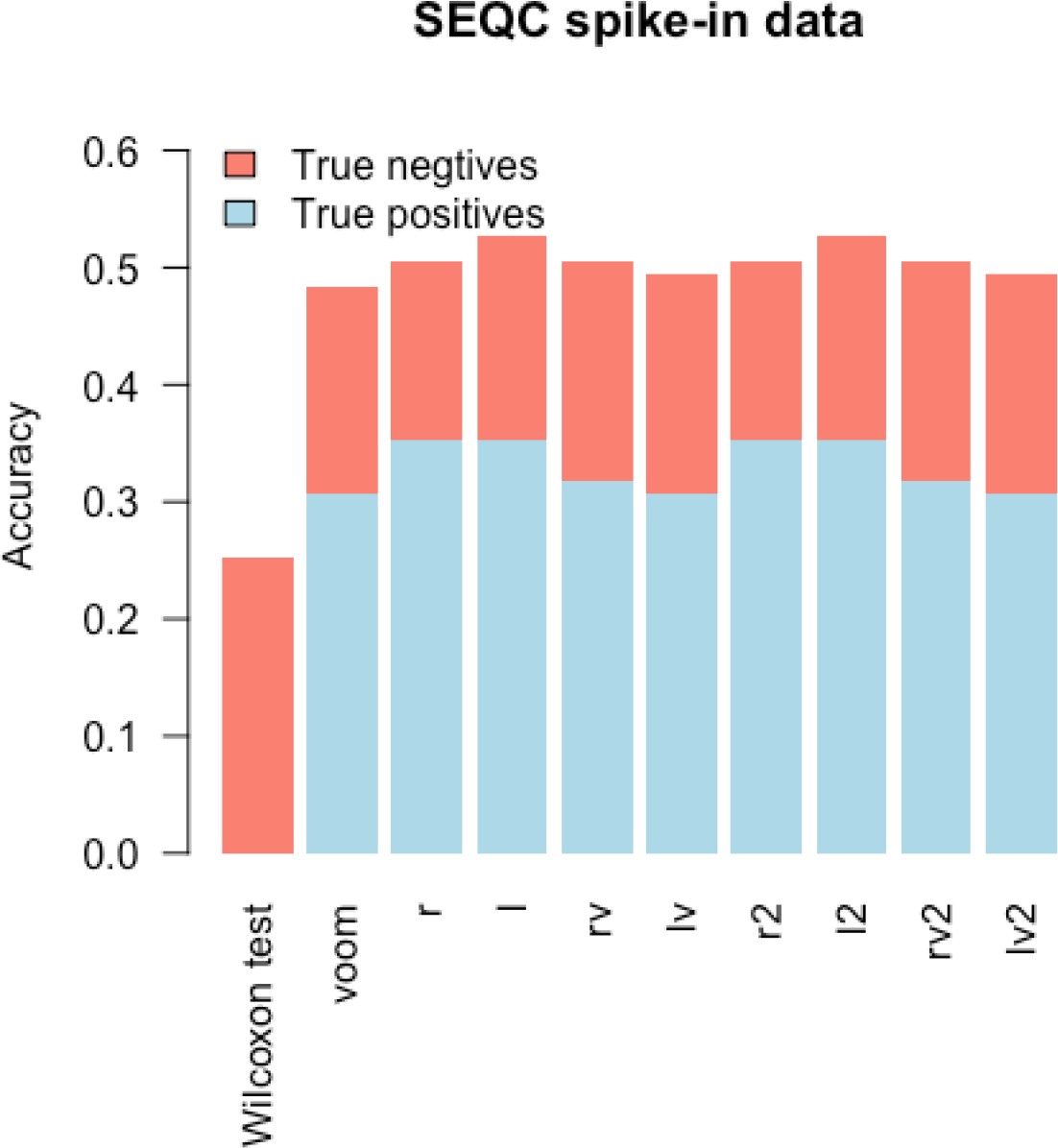
Accuracies obtained based on the SEQC spike-in data. The accuracy (=true positives + true negatives) is plotted for each of the 10 methods. Limma with the *l2* and *l* transformations had the highest accuracies (0.53).

GSE95587 is an RNA-seq dataset for Alzheimer’s disease that has relatively large sample size (n=117). We used all 117 samples (84 Alzheimer samples and 33 age-matched normal controls) and conducted 100 random partitions of the 117 samples. In each random partition, we randomly split the 117 samples into roughly 2 equal sets: discovery set and validation set. We then calculated the proportion of validated DE genes for each of the 10 methods. Figure 7 showed that limma with *l* and *l2* had higher median validation proportion than limma with *Wilcoxon* and limma with other transformations in two analyses. We also observed that limma with the proposed 8 transformations had larger median proportions of validation than or similar proportions to limma with *voom* when sample size is larger.

**Figure 7.**
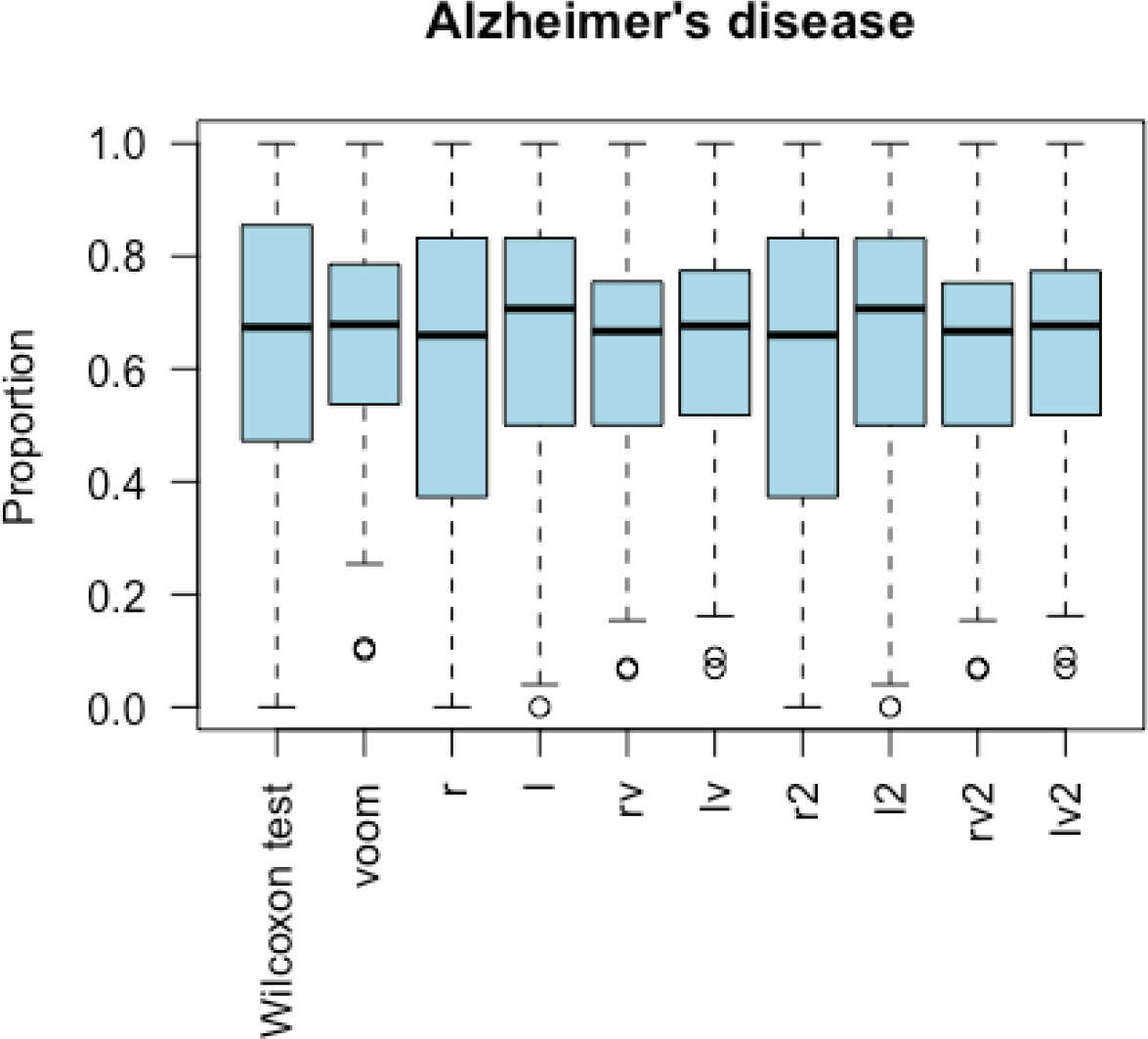
Parallel boxplots of the proportion of validated genes in the analysis of GSE95587. We did 100 randomly splits the 117 samples. In each split, we randomly split the 117 samples into roughly equal two parts: discovery set and validation set. The proportion of the significant DE genes detected in the discovery set and validated in the validation set was recorded for each split. Each boxplot is the summary of the proportions of the 100 proportions. The higher the proportion, the better the performance.

Supplementary Table 2 is a summary of the performances of *Wilcoxon test* and limma with all 9 RNA-seq transformations for real data analyses, we ranked accuracy for two SEQC tests and the proportion of validation for GSE95587, we then obtained the mean ranks to compare the performance. Supplementary Table 2 showed that limma with all 8 proposed transformations had smaller mean ranks than limma with *voom* in three real data analyses.

## Discussion

In this article, we proposed 8 new RNA-seq data transformations to improve the *voom* transformation for RNA-seq data analysis. The simulation results (Supplementary Table 1) showed that overall limma with *rv* had the best performance compared to *Wilcoxon* test, limma with voom, and limma with the other 7 proposed transformations. We also observed that limma with *rv2* had the 2nd best performance when sample size is small (nCases=nControls=3). Limma with the *voom* transformation and limma with the *r2* transformation performed last among the 10 methods compared when nCases=nControls=3. When sample size is medium (nCases=nControls=30 or nCases=nControls=50), limma with *voom*, *rv*, and *rv2* had the best performance. When sample size is large, limma with *r2* had the best performance followed by the *Wilcoxon* test. Limma with the proposed log-transformation-based methods did not perform as well as other methods in the simulation studies. However, limma with the *l* and *l2* transformation performed the best in the real data analyses. The real data analyses also showed that limma with *voom* did not perform as well as the other 9 methods. The inconsistency between the results of simulation studies and those of real data analyses is probably due to the fact that the real data are much more complicated and different than the simulated datasets.

Having sample mean close to sample median for pooled data could not guarantee that for each gene, the sample mean is close to the sample median for cases and for controls, respectively. Also, the empirical distribution of the pooled data after data transformation might still be skewed even if the sample mean is very close to sample median. Hence, robust linear regression might improve the performance of limma after data transformation.

The *voom* transformation was proposed by [12] and has been implemented in the *limma* package. Law et al. (2014)[12] focused on small sample size (nCases=nControls=3) because RNA-seq data were expensive to obtain at that time. Since then the cost of RNA sequencing become lower and lower. Hence, we evaluated the performances of limma with *voom* and limma with the 8 new transformations in scenarios where sample sizes are relatively large (nCases=nControls=100). We also applied the *Wilcoxon* test to the *raw count* data to check if data transformation could perform better than the non-parametric test. Interestingly, the *Wilcoxon* test *without data transformation* performed better than the *limma* analysis based on the 9 data transformations in simulation studies when sample sizes are not too small and sample library sizes are unequal. The analysis of the Alzheimer’s disease dataset also demonstrated this finding. Further investigation is warranted.

In this article, we did not compare *limma* with count-based methods, such as edgeR and DESeq since Law et al. (2014)[12] did the comparison and showed the good performance of the data transformation approach. However, Law et al. (2014) [12] did the comparisons based on datasets with small sample sizes. Moreover, new count-based methods, such as DESeq2, have been proposed since 2014. Hence, it would be a future research to compare the data transformation approach with all available count-based methods using datasets with large sample sizes (e.g., nCases = nControls = 1000).

We observed that none of the 8 transformations could dominate each other, although they performed better than *voom* in most scenarios (Figure 3 and Supplementary Table 1). For example, limma with the *r2* transformation performed best when sample size is large and samples have equal library size, but could not beat limma with *rv* when sample size is small or moderate (e.g., 6, 60 or 100). Another limitation of our study is that in our real data analyses, no independent cohorts are available to do validation. However, we did 100 times of random splits. In each split, we had discovery set and validation set. In future, we will do validation studies when independent validation sets are available. The third limitation of this study is that the 8 proposed data transformations aim to make the sample mean closer to the distribution center after data transformation. However, the transformed distribution might not be close to a normal distribution. Hence, robust linear regression models are needed since ordinary linear regression requires the normality assumption. Future research is warranted on this subject as well.

## Conclusions

In simulation studies, limma with the *rv* transformation performed better than limma with the *voom* transformation for data with small or moderate sample size. For large sample size, limma with the *r2* transformation performed better than limma with the *voom* transformation. In real data analysis, several (*l2*, *l*, *r2*, *r*, *rv*, and *rv2*) of our proposed transformations performed better than *voom*. We hope these novel data transformations could provide investigators more powerful differentially expression analysis using RNA-seq data.

## Materials and methods

### Eight new data transformations

We proposed 8 new data transformations based on the Box-Cox transformation [19]: 4 root transformations (denoted as *r*, *rv*, *r2* and *rv2*, respectively) and 4 log transformations (denoted as *l*, *lv*, *l2* and *lv2*, respectively). The following two properties of the root transformation motivate us to use root transformations: (1) root transformation of zero exists; (2) root transformation could stabilize the variance of count data.

The *r* transformation is defined as:

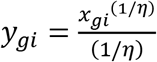

where *x*_*gi*_ is the count of the *g*-th gene for the *i*-th sample. The optimal value for the parameter *η* is to minimize the difference between the sample mean and the sample median of the pooled data:

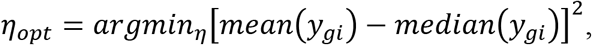

where 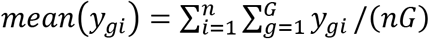 and *median*(*y*_gi_) are the sample mean and sample median of the pool data *y*_*gi*_, i = 1,…, *n*,*g* = 1,…,*G*.

The *rv* transformation is defined as:

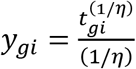

where 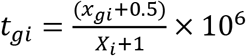, 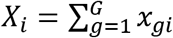

That is, we do the root transformation for the sample-specific counts per million. The optimal value for the parameter *η* is to minimize the difference between the sample mean and the sample median of the pooled data.

The *r2* transformation has the same form as the *r* transformation:

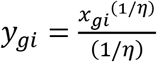

However, the criterion to estimate the optimal value of *η* is different. The optimal value for the parameter *η* is to minimize the difference between the sample mean and the sample median of each sample:

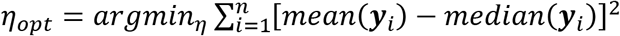

where 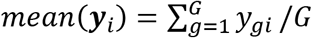 and *median*(**y**_i_)are the sample mean and sample
median of the i-th sample.

The *rv2* transformation is a combination of the *rv* transformation and the *r2* transformation, defined as:

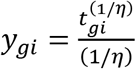

where *t*_*gi*_ is the sample-specific counts per million. The optimal value for the parameter *η* is to minimize the difference between the sample mean and the sample median of each sample.

The *l* transformation is defined as:

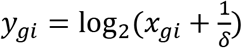

The optimal value for the parameter *δ* is to minimize the difference between the sample mean and the sample median of the pooled data:

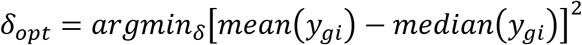

The *lv* transformation is defined as:

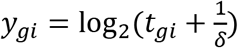

where *t*_*gi*_ is the sample-specific counts per million. The optimal value for the parameter *δ* is to minimize the difference between the sample mean and the sample median of the pooled data.

The *l2* transformation has the same form as the *l* transformation:

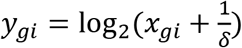

However, the criterion to estimate the optimal value of *δ* is different. The optimal value for the parameter *δ* is to minimize the difference between the sample mean and the sample median of each sample:

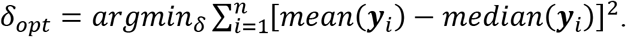

The *lv2* transformation is a combination of the *lv* transformation and the *l2* transformation, defined as:

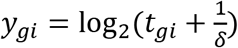

where *t*_*gi*_ is the sample-specific counts per million. The optimal value for the parameter *δ* is to minimize the difference between the sample mean and the sample median of each sample.

### Simulation studies

In each simulation study, we generated 100 datasets. Each dataset contains 10,000 genes, among which 200 genes are differentially expressed between *nCases* cases and *nControls* controls. An inverse chi-square distribution with 40 degrees of freedom was used to generate a modest amount of gene-wise biological variation [12]. We set the number of cases equal to the number of controls (i.e., nCases=nControls).

After data transformation, we used Bioconductor package *limma* to detect differentially expressed genes. We also compared the results based on transformed data with the results of the *Wilcoxon* rank sum test *based on the original counts*.

Law et al. (2014) generated the RNA-seq counts of 10,000 genes for six samples (nCases=nControls=3) with equal or un-equal library sizes in their simulation studies. In our simulation studies, we evaluated the effects of sample size and library size on the performances of the 10 methods (*Wilcoxon*, *voom*, *r*, *l*, *r2*, *l2*, *rv*, *lv*, *rv2*, *lv2*) in detecting differentially expressed genes. We tried four different sample sizes (nCases=nControls=3, nCases=nControls=30, nCases=nControls=50 and nCases=nControls=100) with equal library size and with un-equal library size, respectively.

The criteria to evaluate performance are accuracy, false negative rate (FNR), false discovery rate (FDR), and the difference (DIFF) between the sample mean and the sample median of the pooled expression levels for all samples and all genes. FNR is the percentage of detected non-differentially expressed (NDE) genes among truly differentially expressed (DE) genes. FDR is the percentage of truly NE genes among detected DE genes. Large accuracy and small FNR, FDR, and DIFF indicate good performance.

### Real data analyses

#### SEQC data

Sequencing Quality Control (SEQC) is the third phase of the MAQC project (MAQC-III), aimed at assessing the technical performance of next-generation sequencing platforms by generating benchmark datasets with reference samples [17, 20]. This project provided 6 RNA samples, each sample has 4 replicates, samples A and B were obtained from two well-characterized reference human RNA samples UHR (Universal Human Reference RNA) and HBR (Human Brain Reference RNA). A small amount of Ambion ERCC (External RNA Control Consortium) Spike-in Mix was added into both Sample A and Sample B. Samples C and D were constructed by mixing Samples A and B to known ratios, 3:1 and 1:3, respectively. The pure ERCC Spike-in Mix 1 and 2 were used as Samples E and F. Gene expression levels of Samples A, B, C and D were analyzed by using TaqMan RT-PCR technology.

Our first analysis is based on Samples A (UHR) and B (HBR). The dataset GSE56457 on Gene Expression Omnibus (GEO) website (https://www.ncbi.nlm.nih.gov/geo/query/acc.cgi?acc=GSE56457) provides details about the qRT-PCR data for the SEQC project. We regarded the expression levels of these genes measured by qRT-PCR as the true expression levels. If a gene has mean log2 fold-change (LFC) greater than 2 between two RNA samples in GSE56457, we claimed it as a truly differentially expressed gene. If a gene has mean LFC less than 0.004, we claimed it as a truly non-differentially expressed gene [21]. Based on this criterion, there were 390 DE genes and 457 non-DE genes. We evaluated the performance of the 9 data transformations using SEQC data based on these 847 genes. We applied *limma* to detect differentially expressed genes after data transformation. We also applied the *Wilcoxon* test to detect DE genes based on the raw count data. A gene was estimated as a DE gene if it had FDR-adjusted p-value < 0.05. We then calculated the proportion of agreement (i.e., accuracy) between the true gene significance of the 847 genes and the estimated gene significance.

#### SEQC spike-in (ERCC)

We downloaded ERCC data from http://bioinf.wehi.edu.au/voom/ and did similar analysis based on Samples E and F, which are the ERCC RNA Spike-In Mixes, providing a set of external RNA controls that enable performance assessment of a variety of technology platforms used for gene expression experiments. These 8 samples (4 from Samples E and 4 from Samples F) are pre-formulated sets of 92 poly adenylated genes from the ERCC plasmid reference library, three quarters of the genes were truly DE and the remaining quarter were not. The genes are traceable through the manufacturing process to the NIST plasmid reference material. This dataset provided concentrations of the two mixes, the log2 fold change of concentration can be used for determining if a gene is DE. The analysis procedure of spike-in data is consistent with SEQC data. We calculated the accuracy to compare the transformation methods performance.

#### Neurodegenerative disease RNA samples

GSE95587 (https://www.ncbi.nlm.nih.gov/geo/query/acc.cgi?acc=GSE95587) is an RNA-seq dataset obtained from fusiform gyrus tissue sections of autopsy-confirmed Alzheimer’s cases and neurologically age-matched normal controls. The matching information was not provided in GSE95587. We downloaded the RNA-seq raw data and annotations from Gene Expression Omnibus (GEO, https://www.ncbi.nlm.nih.gov/geo). The dataset contains 117 samples, 84 of which are from Alzheimer’s cases (ADs) and 33 of which are controls (CONs). We randomly split the 117 samples into two roughly equal parts: a discovery set and a validation set. The discovery set has 42 ADs and 17 CONs.

The validation set has 42 ADs and 16 CONs. We then applied *limma* after data transformations to the discovery set and the validation set to detect differentially expressed (DE) genes. For the discovery set, we claimed a gene is DE if its FDR-adjusted p-value < 0.05. For the validation set, we claimed a gene is validated DE if it had a raw p-value < 0.05 in the validation set and it had FDR-adjusted p-value < 0.05 in the discovery set. We repeated the above split-validation procedure 100 times. For each of the 10 methods, we calculated the proportion of the validated DE genes among the DE genes detected in the discovery set. The higher the proportion is, the better performance, the method is.

For the *voom* transformation in all real data analyses in this article, we followed Law et al. (2014) by first applying TMM scale-normalization[18] and quantile normalization before applying for the *voom* transformation.

## Declarations

### Ethics approval and consent to participate

Not applicable. The real data sets we used are downloaded from public data repositories.

### Consent for publication

Not applicable. The real data sets we used are downloaded from public data repositories.

### Availability of data and material

The SEQC data can be downloaded from Gene Expression Omnibus (GEO) with accession number SE56457 (https://www.ncbi.nlm.nih.gov/geo/query/acc.cgi?acc=GSE56457). The SEQC spike-in (ERCC) data can be downloaded from http://bioinf.wehi.edu.au/voom/. The neurodegenerative disease data can be downloaded from GEO with accession number GSE95587 (https://www.ncbi.nlm.nih.gov/geo/query/acc.cgi?acc=GSE95587).

### Authors’ contributions

WQ initiated this work. ZZ and WQ developed the method and wrote the manuscript. ZZ implemented the method and performed the analyses. MS helped clarify the methods.

DY, CH, STW helped interpret the results and manuscript writing. All authors read and approved the final manuscript.

#### Acknowledgements

Not applicable

